# Monitoring fungicide resistance frequencies – a case study of barley net blotch

**DOI:** 10.1101/2025.01.16.633467

**Authors:** N. L. Knight, K. C. Adhikari, W. J. Mair, K. Dodhia, K. G. Zulak, F. J. Lopez-Ruiz

## Abstract

Decreased sensitivity to fungicides impacts the effectiveness of fungicide applications for managing plant disease. Knowledge of the frequency of decreased sensitivity in field populations is critical for evaluating risks for disease control. This study has applied a droplet digital PCR detection approach to assess pathogen populations and quantify the frequencies of alleles associated with decreased sensitivity to either demethylation inhibitor or succinate dehydrogenase inhibitor fungicides in *Pyrenophora teres* causing net blotch on barley in Western Australia. *Pyrenophora teres* f. *maculata* was the most frequent form of the pathogen in the sampled region. Frequencies of decreased fungicide sensitivity alleles varied, being as great as 92% for the PtTi insertion in the CYP51A promoter, 88% for C-S75 and 28% for L489-2. Impacts of cultivar selection and weed hosts on the presence and survival of decreased fungicide sensitivity were observed. Determining the dynamics of alleles within different field populations of *P. teres* provides an important perspective on the impact of fungicides, the fitness associated with decreased fungicide sensitivity alleles and the susceptibility of barley cultivars.

## Introduction

The effectiveness of fungicides is an important concept in plant disease control, and the emergence of decreased fungicide sensitivity in fungal populations adds another level of complexity for disease management. Disease control strategies prioritise planting the least susceptible varieties to reduce disease impacts and integrated disease management techniques to reduce inoculum in the environment. Applications of fungicides are suggested to occur when necessary, with rotations or mixtures of fungicide modes of action recommended to reduce selection pressure for decreased sensitivity (Corkley et al. 2022; Ireland et al. 2021). However, once decreased sensitivity has emerged in a pathogen population, the effectiveness and choice of fungicides becomes less certain. This suggests that knowledge of the frequencyof decreased fungicide sensitivity in the pathogen population is required to develop informed fungicide management strategies.

As reports of decreased fungicide sensitivity in plant pathogens become more prevalent (Corkley et al. 2022), monitoring field environments has increased in importance, with rapid and sensitive tools required to provide growers with information that can be incorporated into control strategies. Decreased fungicide sensitivity is traditionally confirmed through pathogen isolation and phenotyping tests on fungicide amended media (Ishii and Hollomon 2015). While fundamental, there are practical limitations for the number of samples which can be assessed and the time required for isolating and growing cultures (Brent and Hollomon 2007). Genotyping has emerged as a more rapid tool for detecting decreased fungicide sensitivity (Ishii and Hollomon 2015). This relies on the identification of DNA sequences associated with decreased fungicide sensitivity and the development of tests to specifically detect those sequences. These tests may be used on pure fungal DNA, or on mixed DNA extracted directly from affected plants or environmental sources. This approach also moves beyond the ability to just detect the presence of decreased fungicide sensitivity and towards a more comprehensive view of the frequency of associated alleles in a population.

A range of pathogens have been genetically characterised for decreased fungicide sensitivity, with allele-specific PCR for the detection of both the sensitive and decreased sensitive alleles being deployed across PCR platforms to report frequencies of relevant alleles in pathogen populations. Real-time quantitative PCR has been a popular platform for fungicide resistance detection (Capote et al. 2012; Ma and Michailides 2005), with relative quantification of each allele allowing frequency determination (Camiletti et al. 2022; Hellin et al. 2020). Digital PCR, an emerging technology for fungicide sensitivity allele frequency detection, can use the same allele-specific PCR tests and has the advantage of not requiring a standard curve for relative quantification (The dMIQE Group and Huggett 2020). Digital PCR has been used to quantify two *Cyp51* mutations in *Blumeria graminis* f. sp. *hordei* (Zulak et al. 2018), the G143A mutation in *Erysiphe necator* (Miles et al. 2020), *Blumeria graminis* f. sp. *tritici* (Dodhia et al. 2021) and *Zymoseptoria tritici* (Battistini et al. 2022), and a *Cyp51* mutation in *Pyrenophora teres* f. *teres* (Hodgson et al. 2022; Hodgson et al. 2024a; Hodgson et al. 2024b). The ability of these tools to assess multiple alleles in pathogen populations offers the ability to determine current frequencies of each allele in field samples and the potential to monitor changes in response to field management, such as fungicide applications.

*Pyrenophora teres* f. *maculata* and *P. teres* f. *teres*, causing spot form and net form net blotch of barley, respectively, have been phenotypically and genotypically characterised for decreased sensitivity to demethylation inhibitor (DMI; FRAC group 3) and succinate dehydrogenase inhibitor (SDHI; FRAC group 7) fungicides (Campbell and Crous 2002; Ellwood et al. 2019; Mair et al. 2016; Mair et al. 2020; Mair et al. 2023; Rehfus et al. 2016; Sheridan and Grbavac 1985; Sheridan et al. 1985). A workflow comprised of pathogen detection and allele-specific PCR tests for 23 fungicide-sensitivity associated targets has also been reported, making this pathosystem an ideal choice for allele frequency characterisation (Knight et al. 2024). This study aimed to use droplet digital PCR to determine the frequency of alleles associated with decreased sensitivity to DMI and SDHI fungicides in net blotch samples collected from commercial barley fields using a standardised field sampling procedure.

## Materials and methods

### Field sampling

In September 2021, twenty barley fields were sampled for net blotch in Western Australia (Table 1). Sampling was performed at 12 points in total across each field, consisting of three parallel zig-zag transects 50 m apart, each with four points separated by 20 m, covering an area of approximately 0.48 ha. At least ten symptomatic leaves were collected at each point, with a preference for the top three leaves. Barley plants ranged from growth stage 70 to 85 (Zadoks et al. 1974). Two pairs of fields were directly adjacent and planted with two barley cultivars (F21_02/F21_03; F21_10/F21_11), while a third pair was less than 500 m apart (F21_14/F21_15), providing the ability to contrast net blotch severity and allele frequencies between cultivars in the same environment. The cultivars in these fields, RGT Planet and Spartacus CL, are considered susceptible and susceptible to very susceptible, respectively, to *P. teres* f. *maculata*, compared to Maximus CL and Buff, which are rated as moderately susceptible to susceptible and susceptible, respectively (Shackley et al. 2023). Both Maximus CL and Buff represent more recent barley cultivar releases (2021) compared to RGT Planet (2018) and Spartacus CL (2015). Symptomatic leaves of the weed barley grass (*Hordeum glaucum* or *H. leporinum*) were also opportunistically collected from field F21_08. Samples were dried at room temperature.

**Table 1.** Field metadata for barley net blotch sampling performed in 2021.

| Field<sup>a</sup> | Region | Cultivar |
| --- | --- | --- |
| F21_01 | Meckering | Spartacus CL |
| F21_02 | Meckering | RGT Planet |
| F21_03 | Meckering | Maximus CL |
| F21_04 | Doodenanning | Spartacus CL |
| F21_05 | Warding East | Spartacus CL |
| F21_06 | Waeel | Spartacus CL |
| F21_07<sup>b</sup> | Cunderdin | Spartacus CL |
| F21_08 | Youndegin | Spartacus CL |
| F21_09 | Cunderdin | Spartacus CL |
| F21_10 | Nukarni | Spartacus CL |
| F21_11 | Nukarni | Maximus CL |
| F21_12 | Nukarni | Scope CL |
| F21_13 | Meckering | Spartacus CL |
| F21_14 | Quelagetting | Spartacus CL |
| F21_15 | Quelagetting | Buff |
| F21_16 | Watercarrin | Spartacus CL |
| F21_17 | Watercarrin | Spartacus CL |
| F21_18 | Kalannie | Spartacus CL |
| F21_19 | Muresk | Spartacus CL |
| F21_20 | Cunderdin | Spartacus CL |

<sup>a</sup> Sampling in 2021 occurred across three zig-zag transects.

### Droplet digital PCR detection range

The range of detectable DNA concentrations was assessed for the duplex of the form-specific assays described by Knight et al. (2024). DNA was extracted from pure mycelium as described above and a ten-fold dilution series ranging from 10 to 0.0001 ng of DNA was created for *P. teres* f. *maculata* (isolate 16FRG073) and *P. teres* f. *teres* (isolate Ko103). Each concentration was assessed in triplicate in ddPCR (conditions described below). The linear relationship between the log of the mean copy number and the log of the DNA quantity was calculated in Microsoft Excel.

### Droplet digital PCR assessment of allele frequencies in lesions

For each of the fields, a single lesion was excised from five leaves of each sampling point. These were combined across the 12 points to give a total of 60 lesions per field. Tissues were dried at 65 °C overnight and ground for 60 s at 30 Hz (Retsch MM400) in 2 mL tubes with two 3 mm diameter steel ball bearings. DNA was extracted from duplicate 20 mg sub-samples for each field using a BioSprint 15 DNA Plant Kit (Qiagen, Hilden, Germany) according to the manufacturer’s instructions. DNA was eluted into a 200 µL volume and stored at −20 °C.

DNA of each sample was assessed in droplet digital PCR using form-specific and allele-specific assays for targets in the *Cyp51A* promoter region, the *Cyp51A* gene and succinate dehydrogenase (*sdh*) B, C and D subunit genes described by Knight et al. (2024), and the mating type assays described below. Control DNA for each assay was obtained as described by Knight et al. (2024) and included the associated target and non-target alleles, along with a no template control. Each sample and control was assessed in duplicate.

Droplet digital PCR was performed using the QX200 AutoDG Droplet Digital PCR System (Bio-Rad Laboratories) as per the manufacturer’s protocol. Droplet digital PCR reactions were set up in 22 μL volumes containing 11 μL ddPCR Supermix for Probes (No dUTP) or ddPCR EvaGreen Supermix (Bio-Rad Laboratories) and 5 μL of DNA template. For probe-based assays, primer and probe (Integrated DNA Technologies) concentrations were 0.25 μM of each forward and reverse primer and 0.15 μM of probe in uniplex and duplex reactions. In amplitude-based tetraplex reactions, assays 1 and 2 were at the primer/probe concentrations described, while assays 3 and 4 were 0.9 μM of each forward and reverse primer and 0.25 μM of probe. For the EvaGreen-based assay, each forward and reverse primer were 0.1 μM. Thermal cycling for probe-based reactions consisted of an initial denaturation for 10 min at 95 °C followed by 50 cycles of 94 °C for 30 s and the assay specific temperature (Supp. Table S1) for 60 s. Samples were then held at 98 °C for 10 min, before cooling to 4 °C. For EvaGreen-based reactions, thermal cycling consisted of an initial denaturation for 5 min at 95 °C followed by 50 cycles of 95 °C for 30 s, 58 °C for 60 s and 72 °C for 30 s. Samples were then held at 4 °C for 5 min, followed by 90 °C for 5 min, before cooling to 4 °C. The ramp rate was 2 °C/sec. Droplet data was visualised and manually assigned to clusters in the QX Manager Software v. 1.2 (Bio-Rad Laboratories), with data reported as copies/µL. Copy number values less than 1 were considered below the limit of detection and classified as zero values.

### Mating type assay design

Assays for specifically detecting the mating types MAT-1-1-1 and MAT-1-2-1 of *P. teres* f. *maculata* and *P. teres* f. *teres* (Table 2) were designed by aligning sequences available in GenBank and identifying nucleotides specific to each type (Lu et al. 2010). Both primers were designed to align to a single nucleotide polymorphism (SNP) at the 3′ terminus, and a probe was aligned against one of the same SNPs (Supp. Figure S1). DNA of isolates SG1 (*P. teres* f. *maculata*, MAT1-2-1), Ko103 (*P. teres* f. *teres*, MAT1-2-1), 16FRG073 (*P. teres* f. *maculata*, MAT1-1-1) and 15FRG146 (*P. teres* f. *teres*, MAT1-1-1) (A. Martin, personal communication) was used in quantitative real-time PCR (qPCR) to confirm the detection of the target sequences. DNA extraction and qPCR conditions were as described in Knight et al. (2024).

**Table 2.**
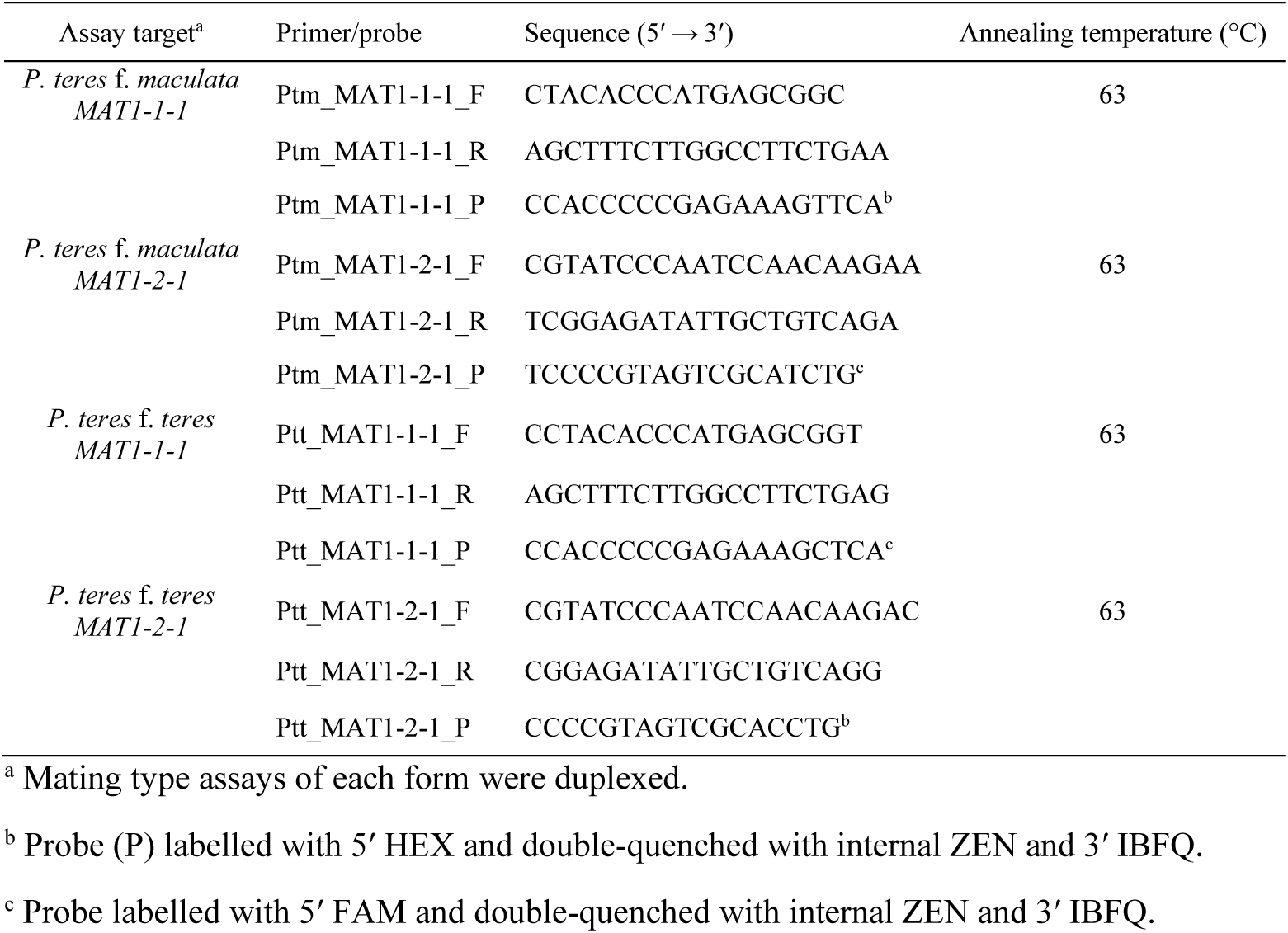
Primer/probe sets for detection of *P. teres* f. *maculata* and *P. teres* f. *teres* mating types.

DNA of each 2021 field sample was assessed using two probe-based duplexed assays for the mating types of each *Pyrenophora* form in ddPCR, as previously described, with an annealing temperature of 61 °C. The mean counts of mating types for each sample were assessed for departure from a 1:1 ratio with a Pearson’s Chi-squared test in RStudio version 2023.06.0.

## Results

### Droplet digital PCR detection range

*Pyrenophora teres* DNA was detectable from 10 to 0.0001 ng, with linear relationships between the log values of DNA quantity and copies/µL reported for both the *Ptm* assay (*y* = 0.93*x* + 2.56, *R^2^*= 0.99) and *Ptt* assay (*y* = 1.01*x* + 2.43, *R^2^*= 0.99). For 10 and 0.0001 ng of DNA this equated to mean values (*n* =3) of 4798.13 and 0.12 copies/µL for the *Ptm* assay and 3208.94 and 0.04 copies/µL for the *Ptt* assay, respectively. Detection of DNA of the non-target form in the 18 dilution series samples was observed for the *Ptm* assay (33%, range of 0 to 0.20 copies/µL, mean of 0.03 copies/µL) and the *Ptt* assay (67%, range of 0 to 0.12 copies/µL, mean of 0.05 copies/µL). These detections occurred across the range of DNA concentrations. A limit of detection for all ddPCR results was set at 1 copy/µL.

### Copy number variation among DNA targets

A comparison of the combined copy numbers for each ddPCR target position, including *Pyrenophora* form, fungicide sensitivity associated and mating type DNA targets, across the sampled region indicated values ranging from 3106 (*Pyrenophora* form) to 3360 (*SdhD*-D+G145) total copies (Fig 1A). This demonstrates that the quantification of target DNA was consistent among the ddPCR assays. Raw allele copy numbers are provided in Supp. Table S2.

**Figure 1.**
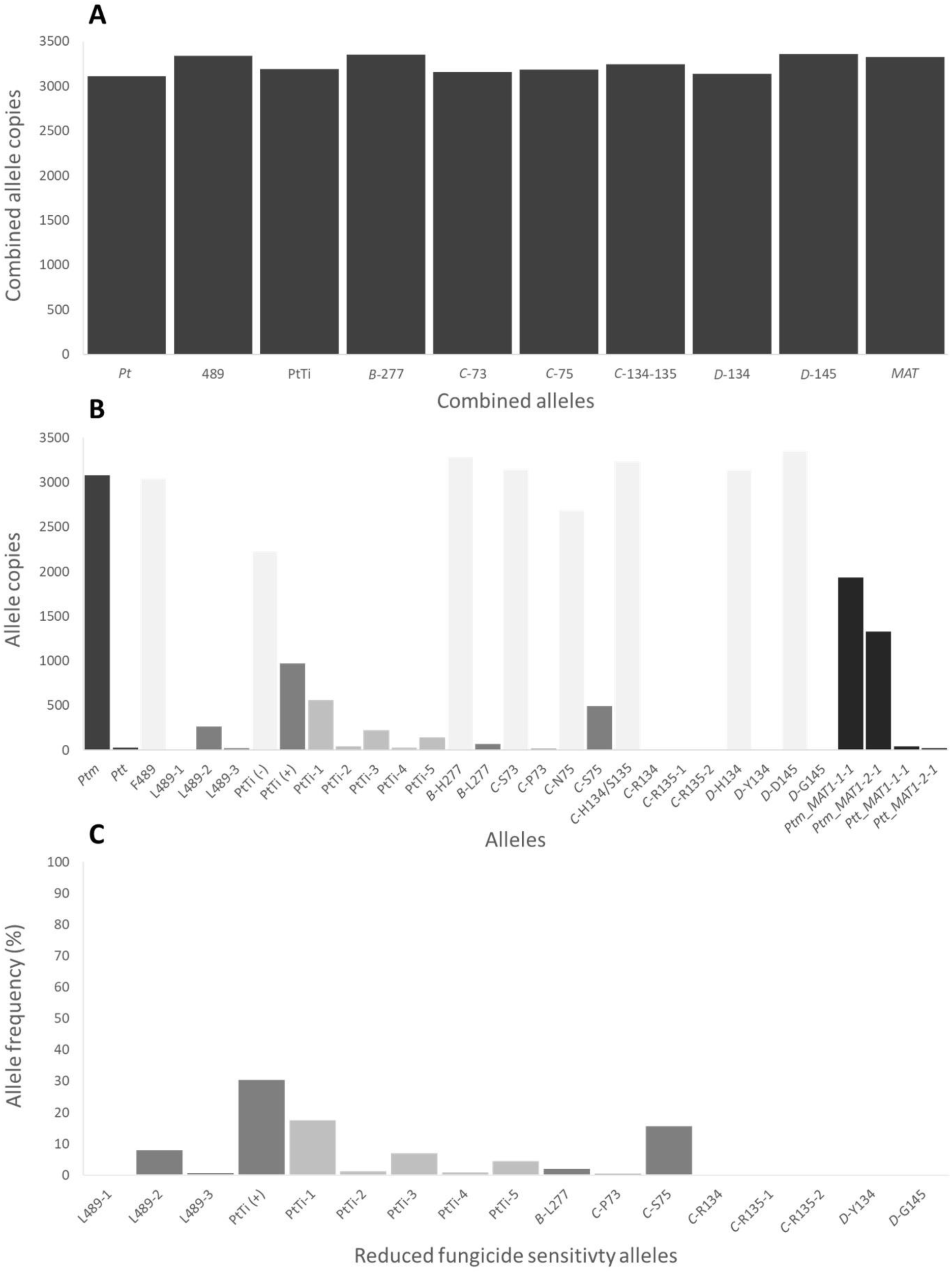
Droplet digital PCR allele copies combined across twenty fields sampled for net blotch in 2021. (A) Combined counts of alleles for *Pyrenophora teres* (*Pt*) forms, position 489 in the *Cyp51A* gene, PtTi insertion position in the *Cyp51A* gene promoter region, six positions in the succinate dehydrogenase sub-units *B*, *C* and *D*, and mating types (*MAT*). (B) Counts of individual alleles at each position, including *P. teres* f. *maculata* (*Ptm*), *P. teres* f. *teres* (*Ptt*), four alleles at the 489 position in the *Cyp51A* gene, the absence [PtTi (-)] and presence [PtTi (+)] of the PtTi insertion in the *Cyp51A* gene promoter region, alleles for six positions in the succinate dehydrogenase sub-units *B*, *C* and *D*, and four mating types (*MAT*). Bar colours indicate alleles associated with fungicide sensitivity (light grey), reduced sensitivity (medium grey) and *Pyrenophora* forms and mating types (dark grey). (C) Frequencies of reduced fungicide sensitivity alleles in the regional population.

### Frequencies of alleles across the sampled region

A combined view of alleles across the sampled region demonstrated the prevalence of *P. teres* f. *maculata*, the total copies of each sensitivity, reduced sensitivity and mating type allele (Fig 1B) and associated frequencies of reduced sensitivity alleles (Fig 1C). For alleles associated with reduced sensitivity to DMI fungicides in the 489 position of *Cyp51A*, the L489-2 allele occurred most often (7.9%). The promoter insertion in *Cyp51A* occurred in 30.4% of the regional population, with the PtTi-1 allele occurring most frequently (17.6%), followed by PtTi-3 (7%), PtTi-5 (4.5%), PtTi-2 (1.3%) and PtTi-4 (0.8%). For the alleles associated with reduced sensitivity to SDHI fungicides, *SdhC*-S75 occurred most frequently (15.6%), followed by *SdhB*-L277 (2%). The remaining alleles occurred at less than 1%.

### Net blotch form and copy numbers within fields

The *P. teres* f. *maculata* allele was detected across all fields sampled in 2021 and ranged from nine to 533 copies/µL (Fig. 2). The *P. teres* f. *teres* allele was detected in three fields and ranged from seven to 12 copies/µL (Fig. 2). This was consistent with visual observations of disease symptoms. Fields varied in fungicide treatments and cropping history (Supp. Table S3).

**Figure 2.**
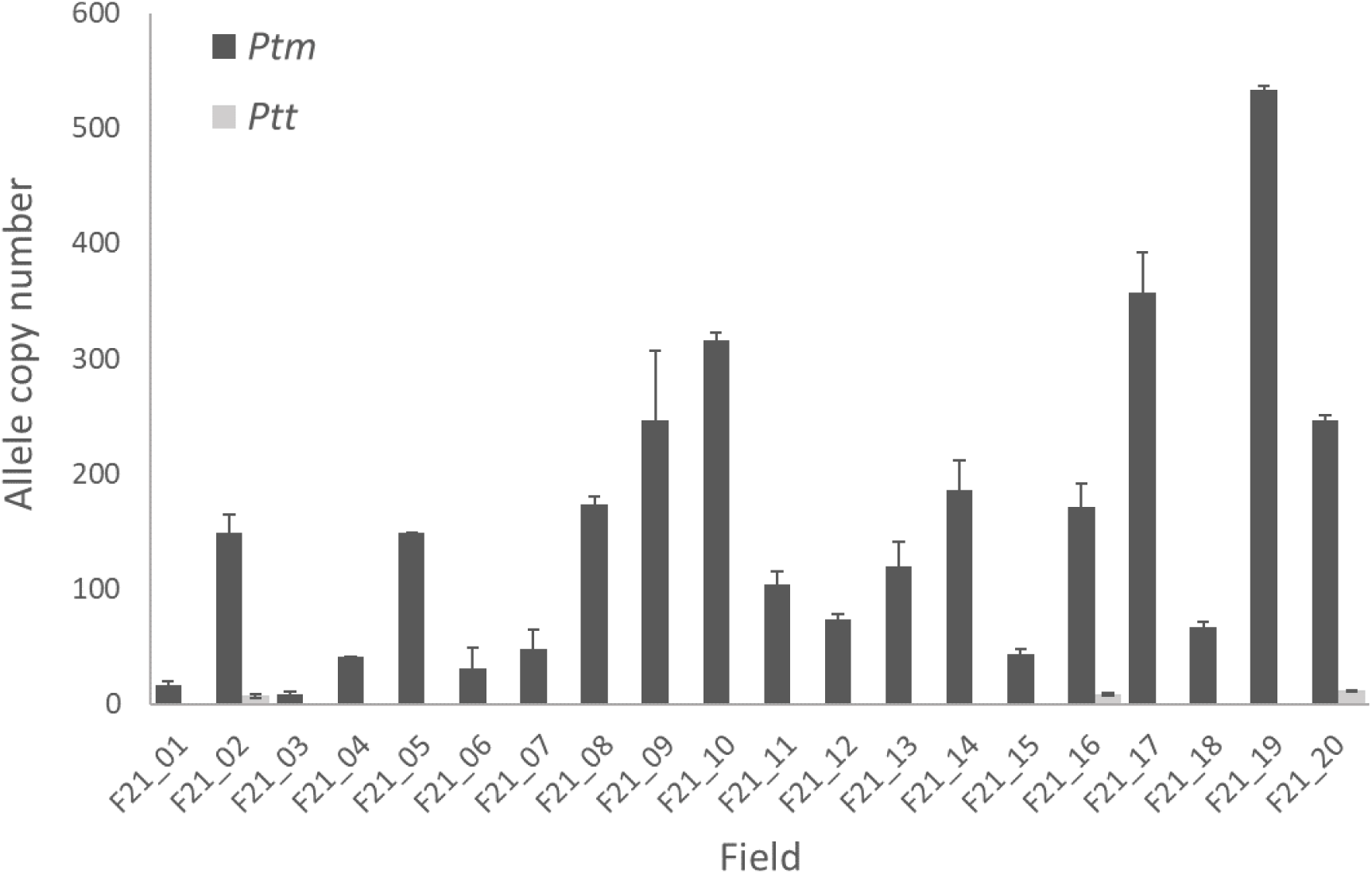
Number of copies of alleles specific to *Pyrenophora teres* f. *maculata* (*Ptm*) or *P. teres* f. *teres* (*Ptt*) in DNA extracted from combined net blotch lesions collected from 20 barley fields in 2021. Bars represent the standard error of the means of two replicate DNA extractions.

### *Cyp51A* allele frequencies within fields

Across individual fields, alleles associated with reduced sensitivity to DMI fungicides in the 489 position of *Cyp51A* were detected in 17 of the 20 fields (ranging from 0 to 31%) (Fig 3A). Allele L489-2 was detected in 17 fields (ranging from 0 to 28.4%), followed by allele L489-3 in seven fields (ranging from 0 to 6.7%) and allele L489-1 in two fields (ranging from 0 to 2%).

**Figure 3.**
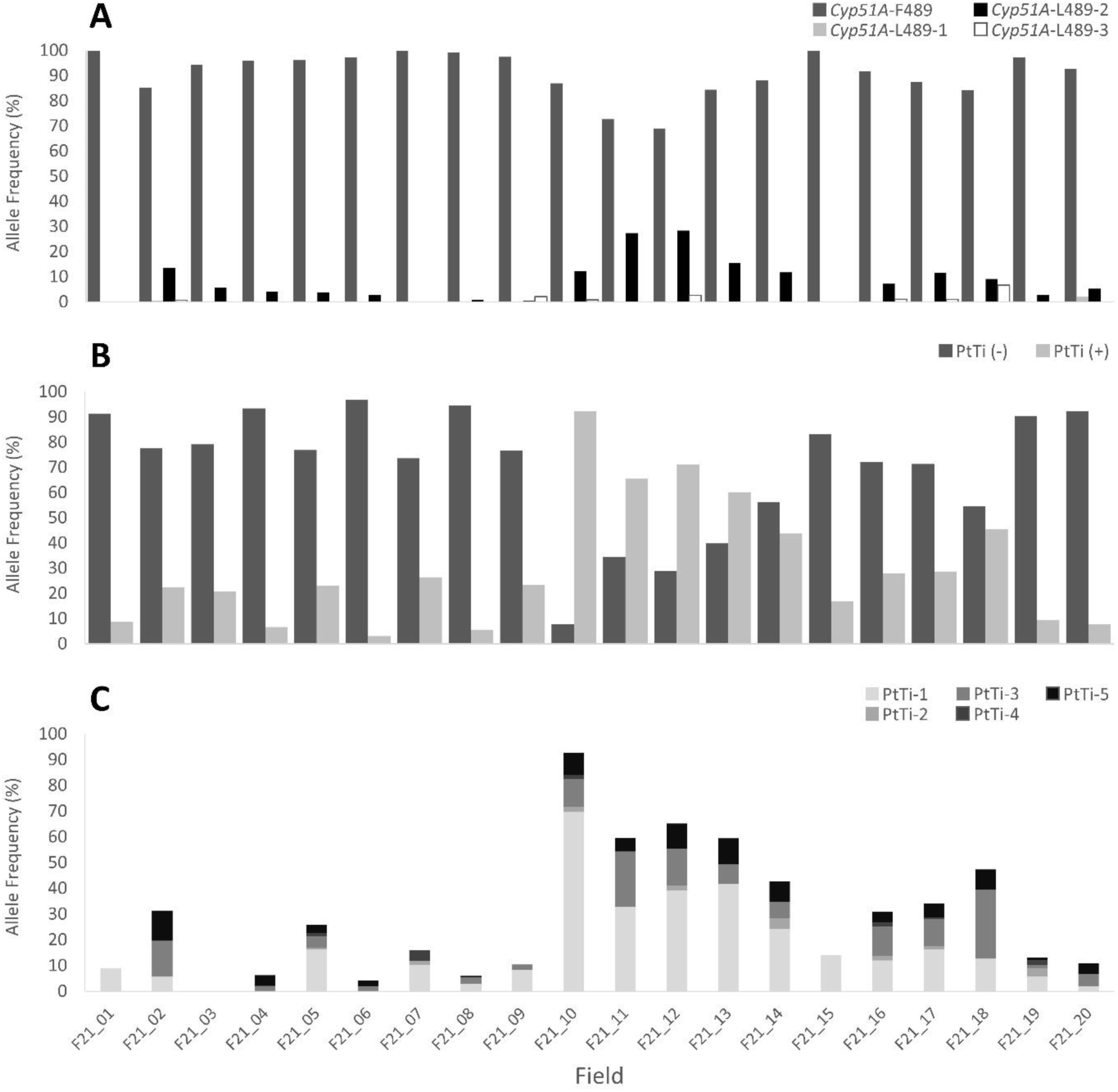
Frequency of alleles in DNA extracted from combined net blotch lesions collected from twenty fields in 2021 associated with sensitivity to demethylation inhibitor fungicides. (A) Frequency of four alleles in the *Cyp51A* gene associated with sensitivity (F489) and reduced sensitivity (L489-1, -2 and -3). (B) Frequency of the absence (-) or presence (+) of the PtTi insertion, associated with reduced sensitivity, in the *Cyp51A* promoter region. (C) Frequency of five positions of the PtTi insertion in the *Cyp51A* promoter region.

The PtTi insertion in the *Cyp51A* promoter was detected in all 20 fields (ranging from 3.1 to 92.3%) (Fig 3B). Allele PtTi-1 was detected in 17 fields (ranging from 1.9 to 69.6%), followed by allele PtTi-3 in 16 fields (ranging from 0.9 to 26.9%), allele PtTi-5 in 15 fields (ranging from 0.5 to 11.7%), allele PtTi-2 in eight fields (ranging from 0.5 to 4.2%) and allele PtTi-4 in six fields (ranging from 0.6 to 4.1%) (Fig 3C). The total copy numbers for the presence of the PtTi insertion compared to the sum of the copy numbers for the specific PtTi alleles were similar within fields, suggesting a low frequency of alternative PtTi insertion positions.

### *Sdh* allele frequencies within fields

Alleles associated with reduced sensitivity to SDHI fungicides were detected in 15 of the 20 fields (ranging from 0.8 to 94.9%) (Fig 4). Allele *SdhC*-S75 was detected in 15 fields (ranging from 0.8 to 88.7%), followed by *SdhB*-L277 in eight fields (ranging from 1.1 to 10.8%), *SdhC*-P73 in four fields (ranging from 0.1 to 4.5%), *SdhC*-R135-2 in four fields (ranging from 0.6 to 4.3%), *SdhD*-G145 in three fields (ranging from 0.4 to 3.6%) and *SdhC*-R134 in two fields (ranging from 0.3 to 0.7%). Alleles *SdhC*-R135-1 and *SdhD*-Y134 were not detected.

**Figure 4.**
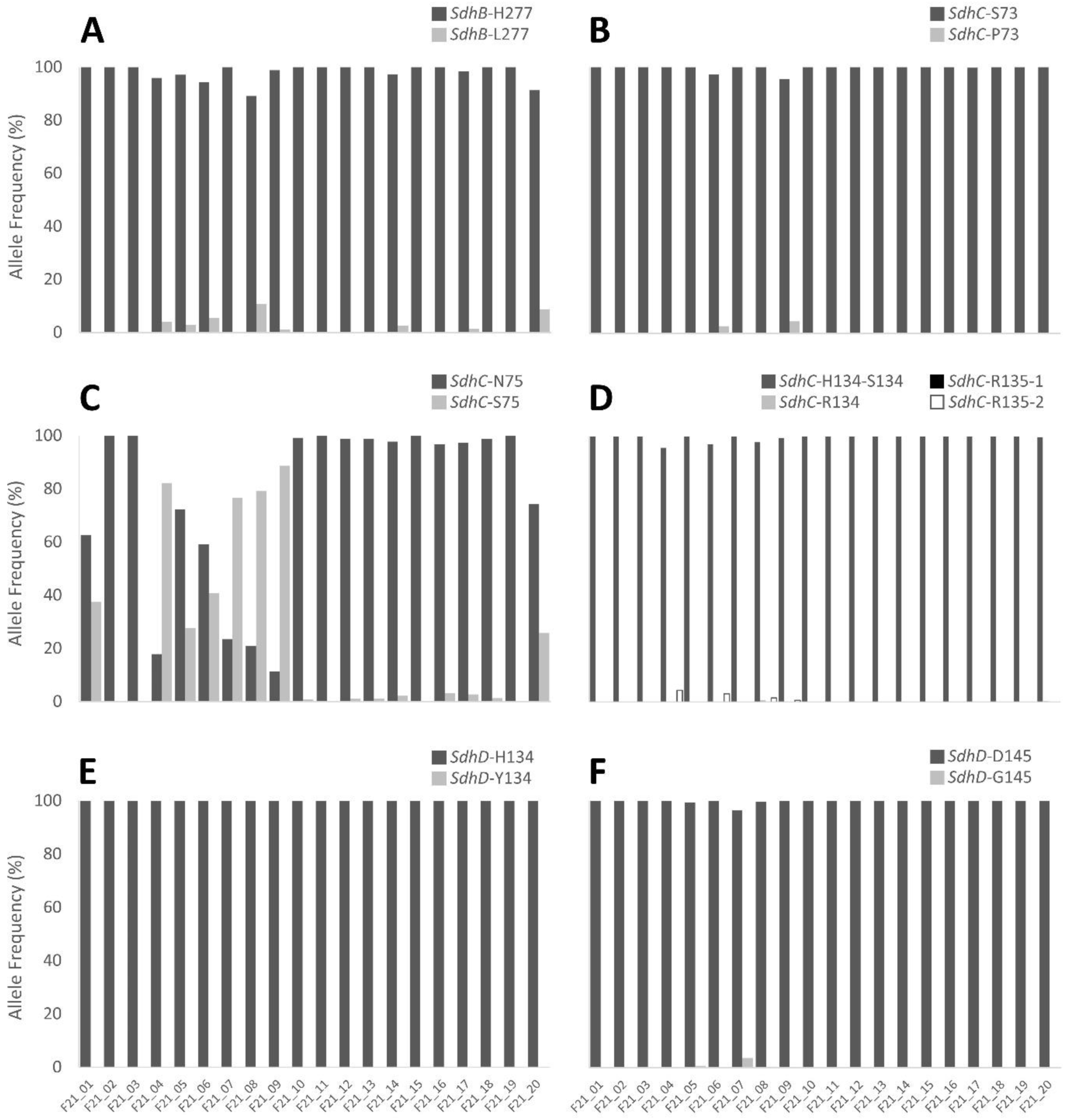
Frequency of alleles in DNA extracted from combined net blotch lesions collected from twenty fields in 2021 associated with sensitivity to succinate dehydrogenase inhibitor fungicides. The dark grey bars correspond to the sensitive allele, and remaining colours correspond to alleles associated with reduced sensitivity.

### Mating type allele frequencies within fields

Both *P. teres* f. *maculata* mating types were detected in all 20 fields (Table 3). There was a significant (*P* < 0.005) departure from a 1:1 ratio (expected under random mating) in ten fields. Of the ten fields, eight were dominated by the *Ptm*_*MAT1-1-1* type. *Pyrenophora teres* f. *teres* mating types were detected in ten fields, with *Ptt*_*MAT1-1-1* detected in all ten fields and *Ptt*_*MAT1-2-1* detected in two fields (Supp. Table S4). These two fields contained both mating types, and a significant (*P* < 0.05) departure from the 1:1 ratio, with *Ptt*_*MAT1-2-1* the dominant type. Analysis of *P. teres* f. *teres* mating types was limited by low copy numbers in each field (ranging from 1 to 13 copies).

**Table 3.** Mating-type copy numbers for *Pyrenophora teres* f. *maculata* determined by droplet digital PCR using DNA extracted from combined net blotch lesions collected from twenty barley fields in 2021. Departure from a 1:1 ratio is indicated by the chi-square (χ2) statistic.

| Field | *MAT1-1-1* | *MAT1-2-1* | $\chi^2$ |
| --- | --- | --- | --- |
| F21_01 | 2 | 17 | 11.84<sup>a</sup> |
| F21_02 | 76 | 104 | 4.36 |
| F21_03 | 7 | 4 | 0.82 |
| F21_04 | 21 | 32 | 2.28 |
| F21_05 | 88 | 84 | 0.09 |
| F21_06 | 17 | 16 | 0.03 |
| F21_07 | 19 | 29 | 2.08 |
| F21_08 | 101 | 91 | 0.52 |
| F21_09 | 161 | 107 | 10.88<sup>a</sup> |
| F21_10 | 300 | 47 | 184.46<sup>a</sup> |
| F21_11 | 79 | 28 | 24.31<sup>a</sup> |
| F21_12 | 45 | 35 | 1.25 |
| F21_13 | 99 | 32 | 34.27<sup>a</sup> |
| F21_14 | 127 | 50 | 33.5<sup>a</sup> |
| F21_15 | 47 | 2 | 41.33<sup>a</sup> |
| F21_16 | 109 | 88 | 2.24 |
| F21_17 | 244 | 107 | 53.47<sup>a</sup> |
| F21_18 | 52 | 21 | 13.16<sup>a</sup> |
| F21_19 | 220 | 296 | 11.19<sup>a</sup> |
| F21_20 | 121 | 137 | 0.99 |

<sup>a</sup> Mating-type copy numbers with significant deviation from a 1:1 ratio ( $P < 0.005$ ). $\chi^2$ value based on one degree of freedom.

### Comparison of net blotch characteristics between cultivars

Adjacent fields F21_02 and F21_03 were planted with cultivars RGT Planet and Maximus CL, respectively. The comparative *P. teres* f. *maculata* allele copy numbers were 149 and nine, respectively, with seven copies of the *P. teres* f. *teres* allele also present in field F21_02 (Fig 2). Alleles associated with reduced sensitivity to DMI fungicides in the 489 position of *Cyp51A* were present at 14.8% and 5.6%, respectively, with all three L489 alleles detected in field F21_02, and only L489-2 detected in F21_03 (Fig 3A). The PtTi insertion in the *Cyp51A* promoter was detected at 22.4% and 20.8%, respectively, with PtTi-1, -3 and -5 detected in F21_02, while only the general presence of the PtTi insertion was detected in F21_03 (Fig 3B and 3C). No specific PtTi allele position was detected. No SDHI reduced sensitivity alleles were detected in either field (Fig 4). Mating types in both fields were not significantly different from a 1:1 ratio (Table 3).

The second pair of adjacent fields, F21_10 and F21_11 were planted with cultivars Spartacus CL and Maximus CL, respectively. The comparative *P. teres* f. *maculata* allele copy numbers were 315 and 104 (Fig 2). Alleles associated with reduced sensitivity to DMI fungicides in the 489 position of *Cyp51A* were present at 13% and 27%, respectively, with L489-2 and -3 detected in field F21_10, and only L489-2 detected in field F21_11 (Fig 3A). The PtTi insertion in the *Cyp51A* promoter was detected at 92.3% and 65.5%, respectively, with PtTi-1, -2, -3, -4 and -5 detected in F21_10, and PtTi-1, -3 and -5 detected in F21_11 (Fig 3B and 3C). The *SdhC*_S75 allele, associated with reduced sensitivity to SDHI fungicides, was detected in F21_10 at 0.8% (Fig 4C). No other SDHI reduced sensitivity alleles were detected in either field. Mating types in both fields were significantly different from a 1:1 ratio (*P* < 0.005), with both skewed toward the *Ptm*_*MAT1-1-1* type (Table 3).

The third pair of fields in close proximity were F21_14 and F21_15, planted with cultivars Spartacus CL and Buff, respectively. The comparative *P. teres* f. *maculata* allele copy numbers were 185 and 43 (Fig 2). Alleles associated with reduced sensitivity to DMI fungicides in the 489 position of *Cyp51A* were present at 11.8% and 0%, respectively, with L489-2 detected in field F21_14 (Fig 3A). The PtTi insertion in the *Cyp51A* promoter was detected at 43.7% and 16.8%, respectively, with PtTi-1, -2, -3 and -5 detected in field F21_14, and PtTi-1 detected in field F21_15 (Fig 3B and 3C). Two alleles associated with reduced sensitivity to SDHI fungicides, *SdhB*_L277 and *SdhC*_S75, were detected in field F21_14, while no SDHI reduced sensitivity alleles were detected in field F21_15 (Fig 4A and 4C). Mating types in both fields were significantly different from a 1:1 ratio (*P* < 0.005), with both skewed toward the *Ptm*_*MAT1-1-1* type (Table 3).

### Characteristics of net blotch on barley grass

DNA extracted from lesions of symptomatic barley grass in field F21_08 contained 39 copies/µL of the *P. teres* f. *maculata* allele. Twenty-eight copies (87.5%) of the *SdhC*-S75 reduced sensitivity allele were detected. Single copies (2.5% frequency) of the reduced sensitivity alleles PtTi-1 (promoter insertion in *Cyp51A*) and *SdhB*-L277 were detected. No other reduced sensitivity alleles were detected. Both mating types *Ptm_MAT1-1-1* (*n* = 26) and *Ptm*_*MAT1-2-1* (*n* = 5) were present, and significantly differed from a 1:1 ratio (*P* < 0.0005, χ2 = 14.2). In contrast, the mating types in the barley net blotch sample from field F21_08 did not significantly differ from a 1:1 ratio.

## Discussion

Decreased sensitivity to fungicides is emerging as a serious threat to disease control, and knowledge of the frequency of decreased sensitivity in the environment is critical to understanding population dynamics and effective management. This study has applied a molecular detection approach to quantify the frequencies of alleles associated with decreased sensitivity to either DMI or SDHI fungicides in *P. teres* causing net blotch on barley. It was found that *P. teres* f. *maculata* was the most frequent form of the pathogen in the sampled region, and that frequencies varied among decreased fungicide sensitivity alleles, being as large as 94% of a population. Determining the dynamics of alleles within different field populations of *P. teres* provides an important perspective on the impact of fungicides, the fitness associated with decreased fungicide sensitivity alleles and the susceptibility of barley cultivars.

The frequency of decreased fungicide sensitivity in pathogen populations can be determined through both traditional isolation and screening approaches or molecular detection of alleles associated with a decreased sensitivity phenotype (Brent and Hollomon 2007; Ishii and Hollomon 2015). While molecular detection methods are founded upon knowledge generated through traditional screening approaches, they provide the advantage of increasing the coverage of a pathogen population and reducing labour. It has been suggested that screening approaches require 300 samples to be assessed to detect decreased fungicide sensitivity at a 1% frequency (Brent and Hollomon 2007), while molecular detection can achieve a similar outcome within a single reaction. The relevance of both approaches is dependent on the technique used to sample the disease and pathogen, which may include undirected ‘spot’ sampling of a symptomatic plant, transect based approaches or air sampling of spores. Each approach has benefits and limitations for interpreting the frequency of decreased sensitivity in a field, however the more intensive strategies for sampling provide immense benefit for comprehensively reporting the field situation (Van der Heyden et al. 2014). The current study applied a practical transect-based sampling approach which was able to be efficiently conducted over multiple fields and provide a snapshot of the pathogen population across an area of approximately 0.48 ha.

Following sample collection, this study used a molecular detection approach to directly assess DNA extracted from a consistent number of leaf lesions for each field. This bypasses the traditional method of fungal isolation and allows the entire population to be assessed. Fungal isolation strategies are limited by the resources required to maintain large numbers of fungal cultures, and have potential bias for selecting isolates which are more competitive under laboratory conditions. In contrast, the molecular detection process can report the presence of all the pathogen types, with a potential limitation of detecting DNA from dead tissue. However, by collecting leaf lesions from upper leaves, it improves the likelihood that these pathogen types have recently been active in causing disease. A ddPCR platform was chosen for molecular detection based on its ability to give independent quantification of PCR targets and the availability of a suite of allele-specific PCR assays for *P. teres* (Knight et al. 2024). The *P. teres* copy numbers in the DNA template from 60 lesions for each field were consistently within the low to optimum range for ddPCR (8 to 540 copies/µL), demonstrating the sample size was effective for detection. There is potential to combine more lesions in future studies to increase the coverage of the population.

Net blotch disease across the twenty sampled fields was primarily caused by *P. teres* f. *maculata*, with a low amount of *P. teres* f. *teres* detected in three fields. This reflects recent disease prevalence in the sampled region (DPIRD-WA 2024). The pathogen copy numbers varied amongst the fields, suggesting impacts due to timing of collection, barley cultivar, fungicide application, field management and the environment. The pathogen copy numbers were compared with the copies of other alleles, which indicated consistent detection of copy numbers across the fields. This was a useful quality control, which indicated that each ddPCR assay was behaving similarly and that there was a low likelihood of missing target DNA due to the presence of alternative alleles. It is possible, however, that novel alleles conferring decreased sensitivity may be present and missed by the allele-specific PCR approach, highlighting the complementary value of a population level sequencing approach (Zulak et al. 2024).

The presence of decreased fungicide sensitivity alleles in every sampled field is of concern for future net blotch management with fungicides. There was a general trend for greater frequencies of decreased fungicide sensitivity alleles based on the history of fungicide use in each field, however clear responses are challenging to discern. When considered as a combined geographic region, the most frequent decreased fungicide sensitivity alleles occurred as the PtTi insertion in the CYP51A promoter (30%, with allele PtTi-1 at 18%), followed by *C*-S75 (16%) and L489-2 (8%). Across the individual fields the frequency of decreased fungicide sensitivity alleles varied, but was as great as 92% for the PtTi insertion in the CYP51A promoter (field F21_10), 88% for *C*-S75 (field F21_09) and 28% for L489-2 (field F21_12). The relatively high frequency of these three alleles may be due to founder effects and/or their relative fitness in the environment (Hawkins and Fraaije 2018). Indeed, the greater frequencies may reflect a combination of fitness traits, including survival of fungicide applications, virulence on barley and fecundity.

Comparing and interpreting frequencies amongst different studies is challenging due to different sampling and measurement approaches, but they provide some context. A stubble-based assessment of commercial fields in Western Australia using a digital PCR platform reported frequencies as great as 42% for the L489-1 decreased DMI fungicide sensitivity allele (Hodgson et al. 2024b). A subsequent study demonstrated an increase in the L489-1 allele from 14.3% to 24.5% following application of the DMI fungicide tebuconazole (Hodgson et al. 2024a). Based on an isolate screening approach, it was determined that a 2018 collection of *P. teres* f. *maculata* from Western Australia had a decreased DMI fungicide sensitivity phenotype frequency of 52%, linked with both L489 and PtTi alleles (Mair et al. 2020). The authors did indicate that a frequency measure was not appropriate based on how isolates were collected, which also highlights the value of the approach in the current study.

In Europe, *P. teres* decreased SDHI fungicide sensitivity allele frequencies were reported as high as 14.5% and 15.6% for allele *C*-R79 in 2013 and 2014, respectively, based on isolate screening (Rehfus et al. 2016). In contrast to the current study, *C*-S75 was detected at 4.5% in 2014. In Estonia, screening of a *P. teres* f. *teres* collection from 2021 indicated 100% of isolates had a decreased DMI fungicide sensitivity allele (L489-3) compared to 78.7% in 2022 (11.5% L489-1 and 67.2% L489-3) (Pütsepp et al. 2024). The PtTi insertion in the CYP51A promoter was also detected in 2022 at 12.1%. For decreased SDHI fungicide sensitivity allele frequencies, *C*-R135 was reported at 30% and 14% in 2021 and 2022, respectively, with the 2022 collection also exhibiting the *C*-R134 allele at 31%. An allele associated with decreased quinone outside inhibitor (QoI) fungicide sensitivity, L129, was also reported at 30% and 12% in the 2021 and 2022 collections, respectively (Pütsepp et al. 2024). While the relevance of interpreting and comparing these values to those of the current study is debatable, there is reasonable agreement in the range of frequencies being reported.

Frequencies of decreased sensitivity alleles in fields represent valuable information, providing both knowledge of the current population and a baseline to examine changes in frequency. This is essential to move from basic presence/absence reporting to following the dynamics of frequency changes across and within seasons, relating this to field management strategies. The availability of three pairs of barley cultivars in the same fields, or in close proximity, provided the opportunity to observe potential differences in frequencies within similar environmental backgrounds. In particular it allowed the comparison of cultivar effect on a measure of disease severity (pathogen copy numbers) and differences in frequencies of decreased fungicide sensitivity alleles. The trend of the more recently released barley cultivars to have lower pathogen copy numbers than the older cultivars suggests both an impact of host resistance and the adaptation of the *P. teres* f. *maculata* population to the older cultivars. This appears to have impacted the frequency and diversity of decreased fungicide sensitivity alleles, with less diversity on the modern cultivars. Impacts of cultivars on pathogen population diversity are limited, with the concept generally linked to cultivar mixtures (Finckh et al. 2000; Papaïx et al. 2018). Differences in diversity may also be linked to an external inoculum source, with a novel allele reported for the modern cultivar in one pair. However, the mating type ratios suggested that the predominant pathogen background was similar for both cultivars in each pair.

*P. teres* inoculum is primarily associated with crop debris, with spores able to move small distances across the landscape (Jordan 1981), likely a few metres to several kilometres (Rieux et al. 2014). However the seed-borne nature of both *P. teres* forms (Khaledi et al. 2024; Louw et al. 1996; Youcef-Benkada et al. 1994) and grower seed-sharing practices are important factors to consider, especially when *P. teres* f. *maculata* populations with decreased fungicide sensitivity have been reported on seed (Louw et al. 1996; Sheridan et al. 1987).

As a sexually reproducing pathogen, the mating types of *P. teres* may be expected to occur in a 1:1 ratio (Akhavan et al. 2015; McLean et al. 2014; Poudel et al. 2019; Rau et al. 2003; Serenius et al. 2007). However, the asexual stage also plays a significant role as an inoculum source from colonised stubble and may affect mating ratios. Based on the ratios observed in the 2021 field populations, ten were in equilibrium (expected for sexual reproduction), but a significant deviation from a 1:1 ratio was observed for the other ten. Such deviations could be due to rapid asexual reproduction and growth of a genotype with greater fitness or introduction of a novel genotype from external sources. The link between mating type ratios and fitness, particularly regarding fungicide impacts, is unclear, however it could be considered for future studies. To the best of the authors knowledge, this is the first investigation of mating type ratios using combined lesions and ddPCR. The approach provides an indication of the amount of each mating type in the field population, and differs from conventional mating type ratio assessments which are based on collections of single-spored isolates.

*Pyrenophora teres* is able to infect and survive on barley grass (*Hordeum leporinum*), which occurs as a weed in and around barley fields (Khan 1973). Pathogen survival on alternative hosts is a risk for effective disease management, however there is conflicting information regarding the infectivity of *P. teres* from barley grass on cultivated barley (Dahanayaka et al. 2024). It has been reported that isolates from barley grass do not infect barley cultivars (Khan 1973) or cause minimal disease responses and are genetically distinct from barley isolates (Linde and Smith 2019). However, a separate investigation has reported that isolates of both forms from barley grass were not host specific, infecting both barley grass and barley cultivars, while also exhibiting genetic divergence and the ability to sexually recombine (Dahanayaka et al. 2024). In the current study, *P. teres* DNA collected from symptomatic barley grass indicated that *P. teres* f. *maculata* was present, and contained alleles associated with decreased sensitivity to both DMI and SDHI fungicides. As the SDHI fungicide was only applied to barley seeds, it also suggests that barley populations can cross-infect barley grass. This demonstrates that there is a reservoir of decreased fungicide sensitivity alleles which can be harboured in field environments on an alternative *P. teres* host, with the potential to recombine with *P. teres* on cultivated barley.

While this study has demonstrated the application of ddPCR to provide evidence of the presence and frequency of a range of alleles associated with decreased fungicide sensitivity in field environments, this information is simply a baseline. Approaches which indicate frequencies and disease severity should be used to provide critical insights into the effectiveness of practices recommended to manage decreased fungicide sensitivity (Bosch et al. 2014; Vestergård et al. 2023). While challenging, tactics to disrupt fungicide resistance in *P. teres* populations need to be demonstrated in commercial situations, providing confidence to growers that this issue can be managed.

## Acknowledgements

Field identification and access was kindly provided by Dan Taylor (DKT Rural Agencies), Dani Whyte (Braeleigh Consulting) and local growers. We thank Dr Chala Turo and students of AGRI3002 – Integrated Pest Management (Curtin University) for assistance during field sampling. This study was conducted at the Centre for Crop and Disease Management, a joint initiative of Curtin University and the Grains Research and Development Corporation (research grant CUR00023).

