## Supplementary material for "Monitoring fungicide resistance frequencies – a case study of barley net blotch": Supp

### Supplementary Tables and Figures

Supplementary Table 1. Primer/probe sets and annealing temperatures for detection of *P. teres* f. *maculata*, *P. teres* f. *teres* and fungicide sensitivity related genotypes in droplet digital PCR.

| Assay target[a] | Primer/probe[b] | Sequence (5′ → 3′)[c] | Annealing temperature (°C) |
|---|---|---|---|
| *P. teres* f. *maculata* | Ptm_r5ID12_F3.2 | GTTGGCTGACTTGTGTAGACAC | 60 |
| | Ptm_r5ID12_B3_R | AAACGCCCTTTAGCAGTCTT | |
| | Ptm_r5ID12_P | CGCTTGCGCGCATGTTCATT[d] | |
| *P. teres* f. *teres* | Ptt_R1ID1_F3 | GACGCGGCTAATGTCGTAA | 60 |
| | Ptt_R1ID1_B3_R | GGCGGTAACAGCACCAAG | |
| | Ptt_R1ID1_P | AATCGGCGGACCGTCAGGAT[e] | |
| *Cyp51A* promoter[h] | CYP51A_promoter_F | GCCACCCTGACGCTAAGAA | 58 |
| | CYP51A_promoter_R | GGAGAGATGGGCGTAGAACA | |
| | CYP51A_promoter_P | CACCTCACGTGACTTAGGGTCACCTG | |
| PtTi insertion | PtTi-1_F | AGAACTGTGTCCAAAAGTAGAGTGTC | 61 |
| | PtTi-1_P | TGTCCTACATCCGACATAGAGAACCATTTC[f] | |
| | PtTi-2_F | TCATCCTATCGACTTGCTTTATGTC | |
| | PtTi-2_P | TACCTCTGTCCTACATCCGACACTTTACTG[g] | |
| | PtTi-3_F | CTAAGAACTGTGTCCAAAAGTAGATGTC | |
| | PtTi-3_P | TGTCCTACATCTGACAGTAGAGAACCATTTC[f] | |
| | PtTi-4_F | TCCAAAAGTAGAGAACCATTTTGTC | |
| | PtTi-4_P | TCCTACATCCGACACATTTCACCCTAT[g] | |
| | PtTi-5_F | ACCCTGACGCTAAGAACTTGTC | |
| | PtTi-5_P | TCCTACATCTGACAGAACTGTGTCCAAA[f] | |
| *Cyp51A* - F489L | CYP51_F489_mm2L_F | GTACCGACTATAGCACCAAG**T**T+**C** | 60 |
| | CYP51_F-L489_R | GCCTCTCCCAGCAAATCT | |
| | CYP51_F489_mm1_P | CTCTAGGGGGCGCGA+**G**A**A**C[f] | |
| | CYP51_L489-1_mm2L_F | GTACCGACTATAGCACCAAGTT+**A** | |
| | CYP51_L489-1_mm1_P | CTCTAGGGGGCGCGA+**T**AAC[f] | |
| | CYP51_L489-2_mm2L_F | GTACCGACTATAGCACCAAGTT+**G** | |
| | CYP51_L489-2_mm1_P | CTCTAGGGGGCGCGA+**C**AAC[g] | |
| | CYP51_L489-3_mm2L_F | GTACCGACTATAGCACAATG+**C** | |
| | CYP51_L489-3_mm1_P | CTAGGGGGCGCGAGA+**G**CAT[g] | |
| *SdhB* - H277L | SDHB_H-L277_F | GCGAGACGAGAAGAAGG | 56 |
| | SDHB_H277_mm1_R | GAGCAGTTGAGAATGGAG**T** | |
| | SDHB_H277_mm1_P | CATGAGCTTATAGCGATGCC+**A**CAC[f] | |
| | SDHB_L277_L_R | GAGCAGTTGAGAATGGTG+**A** | |
| | SDHB_L277_mm1_P | CATGAGCTTATAGCGATGCC+**T**CAC[g] | |
| *SdhC* - S73P | SDHC_S73_mm2L_F | CACCTGGTTGGCGTCC+**T** | 54 |
| | SDHC_S-P73_R | AGTGTAGGGAGCGACAAGGTA | |
| | SDHC_S73_mm1_P | AGTGATCCGGTTGAGCG+**A**GGA[f] | |
| | SDHC_P73_L_F | CCTGGTTGGCCTCC+**C** | |
| | SDHC_P73_mm1_P | ATGCGCTTGAGCG+**G**GGA[g] | |

| | | | |
|---|---|---|---|
| *SdhC* - N75S | SDHC_N75_mm2_F | GTTGGCCTCCTCGATCA**A** | 60 |
| | SDHC_N-S75_R | GTGTAGGGAGCGACAAGGTA | |
| | SDHC_N75_mm1_P | AGAACCATACCAGTGATGCGG+**T**TGA[f] | |
| | SDHC_S75_mm1_F | GTTGGCCTCCTCGCTTA**G** | |
| | SDHC_S75_mm1_P | AGAACCATACCAGTGATGCGG+**C**TGA[g] | |
| *SdhC* - H134R/S135R | SDHC_H-R134_S-R135_F | CCTACACTGGATGGCACCTC | 60 |
| | SDHC_H134-S135_mm2L_R | CCTCAATCCGTAAAGGCTG+**T** | |
| | SDHC_H134-S135_mm1_P | TTCCCGTTCTTCTTTC+**A**CAG+**C**CTT[g] | |
| | SDHC_R134_mm2L_R | CTCAATCCGTTAACGCTG+**C** | 54 |
| | SDHC_R134_mm1_P | CGTTCCGCTTCTTCTTTC+**G**CAG[f] | |
| | SDHC_R135-1_mm1_R | CAAATGCCTCAATCCGTTATG**T** | 60 |
| | SDHC_R135-1_mm1_P | TTCCGCTTCTTCTTTCACAG+**A**CTT[f] | |
| | SDHC_R135-2_mm1_R | CAAATGCCTCAATCCGTTATG**C** | 64 |
| | SDHC_R135-2_mm1-3_P | TTCCCGTTCTTCTTTCACA+G+**G**+CTT[f] | |
| *SdhD* - H134Y | SDHD_H134_mm2_F | TGCGCTCTTCTGGTCGTC**C** | 60 |
| | SDHD_H-Y134_R | TCGACGATGCATGATCTGGG | |
| | SDHD_H134_mm1_P | ACTCAAAAGCAATGTGTGAGT+**G**GAC[f] | |
| | SDHD_Y134_mm2_F | TGCGCTCTTCTGGTCGTC**T** | |
| | SDHD_Y134_mm1_P | ACTCAAAAGCAATGTGTGAGT+**A**GAC[g] | |
| *SdhD* - D145G | SDHD_D145_L_F | CCAGATCATGCATCGTCG+**A** | 56 |
| | SDHD_D-G145_R | GAGAACAACGGTTCCAGCAC | |
| | SDHD_D145_mm1_P | CGCGCTTCTTGCGGAAATAA+**T**CGA[f] | |
| | SDHD_G145_L_F | CAGATCATGCATCGTCG+**G** | |
| | SDHD_G145_mm1_P | CGCGCTTCTTGCGGAAATAA+**C**CGA[g] | |
| *P. teres* f. *maculata MAT1-1-1* | Ptm_MAT-1-1-1_F | CTACACCCATGAGCGG**C** | 61 |
| | Ptm_MAT-1-1-1_R | AGCTTTCTTGGCCTTCTGA**A** | |
| | Ptm_MAT-1-1-1_P | CCACCCCCGAGAAAGTTCA[d] | |
| *P. teres* f. *maculata MAT1-2-1* | Ptm_MAT-1-2-1_F | CGTATCCCAATCCAACAAGA**A** | 61 |
| | Ptm_MAT-1-2-1_R | TCGGAGATATTGCTGTCAG**A** | |
| | Ptm_MAT-1-2-1_P | TCCCCGTAGTCGCA**T**CTG[e] | |
| *P. teres* f. *teres MAT1-1-1* | Ptt_MAT-1-1-1_F | CCTACACCCATGAGCGG**T** | 61 |
| | Ptt_MAT1-1-1_R | AGCTTTCTTGGCCTTCTGA**G** | |
| | Ptt_MAT1-1-1_P | CCACCCCCGAGAAAG**C**TCA[e] | |
| *P. teres* f. *teres MAT1-2-1* | Ptt_MAT-1-2-1_F | CGTATCCCAATCCAACAAGA**C** | 61 |
| | Ptt_MAT-1-2-1_R | CGGAGATATTGCTGTCAG**G** | |
| | Ptt_MAT-1-2-1_P | CCCCGTAGTCGCA**C**CTG[d] | |

[a] Duplex assays included *P. teres* f. *maculata*+*P. teres* f. *teres*, PtTi-2+PtTi-3, PtTi-4+PtTi-5, F489+L489-2, L489-1+L489-3, *SdhB*-L277+ *SdhC*-S73, *SdhC*-N75+ S75, *SdhD*-H134+ Y134, *SdhD*-D145+ G145, *P. teres* f. *maculata* MAT1-1-1+MAT1-2-1, *P. teres* f. *teres* MAT1-1-1+MAT1-2-1. For 2021 samples, amplitude-based tetraplex assays included F489+L489-2+L489-1+L489-3 and PtTi-2+PtTi-3+PtTi-4+PtTi-5.

[b] Assays for fungicide sensitivity related alleles included a common forward (F) or reverse (R) primer. Each common primer is listed once. Primer CYP51A_promoter_R was used for the PtTi insertion assays.

[c] Locked nucleic acid bases are preceded by a '+'. Deliberate mismatch bases are underlined. Target single nucleotide polymorphisms are in bold.

[d] Probe (P) labelled with 5′ HEX and double-quenched with internal ZEN and 3′ IBFQ.

[e] Probe labelled with 5′ FAM and double-quenched with internal ZEN and 3′ IBFQ.

[f] Probe labelled with 5′ FAM and quenched with 3′ BHQ1 or IBFQ.

[g] Probe labelled with 5′ HEX and quenched with 3′ BHQ1 or IBFQ.

[h] The *Cyp51A* promoter assay was performed with primers only using the ddPCR EvaGreen Supermix to detect both presence and absence of the PtTi insertion or with a probe using the ddPCR Supermix for Probes to detect only the PtTi insertion.

Supplementary Table S2. Allele counts (copies/µL) determined by droplet digital PCR for replicate DNA extractions of tissue samples taken from combined net blotch lesions collected from 20 barley fields in 2021. Values represent the mean of duplicate ddPCR reactions for each sample.

| Field | Sample | Region | Cultivar | Allele copy numbers | | | | | | | | | | | | | | | | | | | | | | | | | | | | | | | |
|---|---|---|---|---|---|---|---|---|---|---|---|---|---|---|---|---|---|---|---|---|---|---|---|---|---|---|---|---|---|---|---|---|---|---|---|
| | | | | *Ptm* | *Ptt* | F489 | L489-1 | L489-2 | L489-3 | PtTi (-)[a] | PtTi (+) | PtTi (+)[b] | PtTi-1 | PtTi-2 | PtTi-3 | PtTi-4 | PtTi-5 | *B*-H277 | *B*-L277 | *C*-S73 | *C*-P73 | *C*-N75 | *C*-S75 | *C*-H134/S135 | *C*-R134 | *C*-R135-1 | *C*-R135-2 | *D*-H134 | *D*-Y134 | *D*-D145 | *D*-G145 | *Ptm_MAT1-1-1* | *Ptm_MAT1-2-1* | *Ptt_MAT1-1-1* | *Ptt_MAT_1-2-1* |
| F21_01 | 1 | Meckering | Spartacus CL | 13 | 0 | 15 | 0 | 0 | 0 | 14 | 2 | 1 | 1 | 0 | 0 | 0 | 0 | 15 | 0 | 16 | 0 | 7 | 4 | 15 | 0 | 0 | 0 | 15 | 0 | 15 | 0 | 2 | 14 | 0 | 0 |
| | 2 | | | 20 | 0 | 24 | 0 | 0 | 0 | 20 | 2 | 2 | 2 | 0 | 0 | 0 | 0 | 22 | 0 | 25 | 0 | 11 | 7 | 22 | 0 | 0 | 0 | 18 | 0 | 21 | 0 | 3 | 20 | 0 | 0 |
| F21_02 | 1 | Meckering | RGT Planet | 133 | 6 | 143 | 0 | 22 | 1 | 128 | 37 | 38 | 9 | 0 | 15 | 0 | 17 | 162 | 0 | 141 | 0 | 149 | 0 | 156 | 0 | 0 | 0 | 153 | 0 | 155 | 0 | 70 | 94 | 7 | 0 |
| | 2 | | | 164 | 9 | 187 | 2 | 31 | 2 | 157 | 46 | 51 | 11 | 0 | 36 | 0 | 26 | 182 | 0 | 163 | 0 | 180 | 0 | 193 | 0 | 0 | 0 | 157 | 0 | 193 | 0 | 83 | 113 | 10 | 0 |
| F21_03 | 1 | Meckering | Maximus CL | 7 | 0 | 7 | 0 | 0 | 0 | 7 | 2 | 2 | 0 | 0 | 0 | 0 | 0 | 9 | 0 | 7 | 0 | 8 | 0 | 6 | 0 | 0 | 0 | 7 | 0 | 8 | 0 | 5 | 3 | 0 | 0 |
| | 2 | | | 10 | 0 | 13 | 0 | 1 | 0 | 10 | 3 | 3 | 0 | 0 | 0 | 0 | 0 | 13 | 0 | 13 | 0 | 12 | 0 | 12 | 0 | 0 | 0 | 11 | 0 | 11 | 0 | 9 | 5 | 0 | 0 |
| F21_04 | 1 | Doodenanning | Spartacus CL | 41 | 0 | 49 | 0 | 2 | 0 | 51 | 3 | 3 | 0 | 0 | 0 | 0 | 2 | 46 | 2 | 44 | 0 | 8 | 28 | 43 | 0 | 0 | 2 | 46 | 0 | 49 | 0 | 20 | 32 | 0 | 0 |
| | 2 | | | 41 | 0 | 47 | 0 | 2 | 0 | 43 | 3 | 2 | 0 | 0 | 2 | 0 | 2 | 44 | 2 | 46 | 0 | 6 | 33 | 39 | 0 | 0 | 2 | 44 | 0 | 43 | 0 | 22 | 32 | 0 | 0 |
| F21_05 | 1 | Warding East | Spartacus CL | 148 | 0 | 157 | 0 | 6 | 0 | 115 | 34 | 40 | 26 | 1 | 6 | 1 | 5 | 155 | 5 | 140 | 0 | 105 | 34 | 139 | 0 | 0 | 0 | 147 | 0 | 157 | 0 | 86 | 76 | 0 | 0 |
| | 2 | | | 149 | 0 | 155 | 0 | 7 | 0 | 123 | 37 | 40 | 25 | 0 | 8 | 3 | 5 | 171 | 4 | 167 | 0 | 104 | 46 | 150 | 0 | 0 | 0 | 154 | 0 | 155 | 2 | 89 | 91 | 4 | 0 |
| F21_06 | 1 | Waeel | Spartacus CL | 12 | 0 | 13 | 0 | 0 | 0 | 13 | 0 | 0 | 0 | 0 | 0 | 0 | 0 | 12 | 0 | 11 | 0 | 6 | 6 | 13 | 0 | 0 | 0 | 12 | 0 | 13 | 0 | 6 | 6 | 0 | 0 |
| | 2 | | | 49 | 0 | 50 | 0 | 2 | 0 | 70 | 3 | 2 | 0 | 0 | 2 | 0 | 2 | 51 | 3 | 63 | 2 | 30 | 19 | 45 | 0 | 0 | 2 | 47 | 0 | 51 | 0 | 28 | 25 | 0 | 0 |
| F21_07 | 1 | Cunderdin | Spartacus CL | 32 | 0 | 33 | 0 | 0 | 0 | 26 | 8 | 8 | 3 | 1 | 0 | 0 | 0 | 31 | 0 | 33 | 0 | 6 | 18 | 34 | 0 | 0 | 0 | 32 | 0 | 32 | 0 | 13 | 18 | 0 | 0 |
| | 2 | | | 65 | 0 | 68 | 0 | 0 | 0 | 46 | 19 | 19 | 7 | 0 | 0 | 4 | 0 | 48 | 0 | 73 | 0 | 14 | 46 | 68 | 0 | 0 | 0 | 68 | 0 | 60 | 3 | 24 | 41 | 0 | 0 |
| F21_08 | 1 | Youndegin | Spartacus CL | 166 | 0 | 178 | 0 | 1 | 0 | 163 | 9 | 9 | 5 | 0 | 4 | 0 | 0 | 168 | 21 | 178 | 0 | 33 | 97 | 177 | 1 | 0 | 2 | 175 | 0 | 181 | 0 | 93 | 81 | 1 | 0 |
| | 2 | | | 180 | 0 | 207 | 0 | 2 | 0 | 184 | 11 | 11 | 6 | 0 | 5 | 0 | 2 | 178 | 21 | 201 | 0 | 35 | 158 | 197 | 2 | 0 | 4 | 172 | 0 | 199 | 2 | 109 | 100 | 4 | 0 |
| F21_09 | 1 | Cunderdin | Spartacus CL | 186 | 0 | 218 | 0 | 2 | 12 | 154 | 41 | 44 | 15 | 0 | 5 | 0 | 0 | 199 | 2 | 174 | 8 | 14 | 107 | 186 | 0 | 0 | 1 | 181 | 0 | 199 | 0 | 115 | 72 | 1 | 1 |
| | 2 | | | 307 | 0 | 318 | 0 | 0 | 0 | 237 | 78 | 90 | 28 | 0 | 5 | 0 | 0 | 345 | 4 | 321 | 15 | 25 | 204 | 333 | 0 | 0 | 2 | 322 | 0 | 339 | 0 | 208 | 142 | 8 | 0 |
| F21_10 | 1 | Nukarni | Spartacus CL | 323 | 0 | 288 | 0 | 41 | 3 | 23 | 284 | 277 | 208 | 11 | 19 | 0 | 22 | 350 | 0 | 325 | 0 | 364 | 3 | 367 | 0 | 0 | 0 | 324 | 0 | 339 | 0 | 331 | 32 | 2 | 0 |
| | 2 | | | 308 | 0 | 288 | 0 | 40 | 3 | 22 | 262 | 271 | 204 | 1 | 45 | 11 | 28 | 357 | 0 | 335 | 0 | 340 | 3 | 349 | 0 | 0 | 0 | 320 | 0 | 330 | 0 | 268 | 61 | 10 | 0 |
| F21_11 | 1 | Nukarni | Maximus CL | 92 | 0 | 78 | 0 | 29 | 0 | 23 | 58 | 51 | 25 | 0 | 22 | 0 | 3 | 98 | 0 | 90 | 0 | 98 | 0 | 94 | 0 | 0 | 0 | 89 | 0 | 93 | 0 | 71 | 21 | 0 | 0 |
| | 2 | | | 116 | 0 | 76 | 0 | 29 | 0 | 41 | 62 | 62 | 35 | 0 | 17 | 0 | 7 | 125 | 0 | 120 | 0 | 123 | 0 | 122 | 0 | 0 | 0 | 119 | 0 | 124 | 0 | 87 | 36 | 0 | 0 |
| F21_12 | 1 | Nukarni | Scope CL | 78 | 0 | 57 | 0 | 22 | 3 | 25 | 61 | 56 | 33 | 2 | 14 | 0 | 6 | 78 | 0 | 66 | 0 | 80 | 2 | 77 | 0 | 0 | 0 | 81 | 0 | 85 | 0 | 49 | 37 | 0 | 0 |
| | 2 | | | 70 | 0 | 43 | 0 | 19 | 1 | 20 | 50 | 49 | 28 | 1 | 8 | 0 | 9 | 72 | 0 | 63 | 0 | 70 | 0 | 66 | 0 | 0 | 0 | 66 | 0 | 71 | 0 | 41 | 32 | 0 | 0 |
| F21_13 | 1 | Meckering | Spartacus CL | 99 | 0 | 83 | 0 | 15 | 0 | 39 | 56 | 60 | 41 | 0 | 13 | 0 | 4 | 105 | 0 | 102 | 0 | 107 | 0 | 99 | 0 | 0 | 0 | 90 | 0 | 107 | 0 | 74 | 23 | 0 | 0 |
| | 2 | | | 141 | 0 | 130 | 0 | 24 | 0 | 57 | 89 | 86 | 59 | 0 | 6 | 0 | 20 | 168 | 0 | 153 | 0 | 161 | 3 | 156 | 0 | 0 | 0 | 135 | 0 | 161 | 0 | 124 | 40 | 2 | 0 |
| F21_14 | 1 | Quelagetting | Spartacus CL | 212 | 0 | 190 | 0 | 24 | 0 | 112 | 89 | 91 | 49 | 9 | 15 | 0 | 13 | 213 | 6 | 187 | 0 | 224 | 5 | 213 | 0 | 0 | 0 | 209 | 0 | 224 | 0 | 147 | 56 | 0 | 0 |
| | 2 | | | 159 | 0 | 143 | 0 | 20 | 0 | 88 | 67 | 70 | 37 | 6 | 8 | 0 | 15 | 158 | 4 | 149 | 0 | 170 | 4 | 157 | 0 | 0 | 0 | 145 | 0 | 168 | 0 | 107 | 44 | 2 | 0 |
| F21_15 | 1 | Quelagetting | Buff | 38 | 0 | 38 | 0 | 0 | 0 | 38 | 7 | 7 | 6 | 0 | 0 | 0 | 0 | 43 | 0 | 36 | 0 | 42 | 0 | 42 | 0 | 0 | 0 | 39 | 0 | 39 | 0 | 42 | 0 | 0 | 0 |
| | 2 | | | 48 | 0 | 50 | 0 | 0 | 0 | 52 | 11 | 10 | 9 | 0 | 0 | 0 | 0 | 55 | 0 | 47 | 0 | 52 | 0 | 52 | 0 | 0 | 0 | 52 | 0 | 55 | 0 | 53 | 3 | 0 | 0 |
| F21_16 | 1 | Watercarrin | Spartacus CL | 192 | 10 | 209 | 0 | 17 | 2 | 153 | 61 | 64 | 27 | 1 | 27 | 6 | 11 | 205 | 0 | 202 | 0 | 197 | 8 | 210 | 0 | 0 | 0 | 201 | 0 | 213 | 0 | 118 | 101 | 5 | 11 |
| | 2 | | | 150 | 7 | 166 | 0 | 12 | 2 | 128 | 48 | 50 | 20 | 6 | 18 | 0 | 5 | 182 | 0 | 160 | 0 | 174 | 4 | 165 | 0 | 0 | 0 | 160 | 0 | 183 | 0 | 101 | 76 | 0 | 8 |
| F21_17 | 1 | Watercarrin | Spartacus CL | 322 | 0 | 302 | 0 | 41 | 4 | 220 | 88 | 103 | 55 | 1 | 32 | 4 | 29 | 346 | 5 | 336 | 2 | 342 | 10 | 325 | 0 | 0 | 0 | 345 | 0 | 358 | 0 | 212 | 115 | 9 | 0 |
| | 2 | | | 392 | 0 | 346 | 0 | 44 | 3 | 264 | 106 | 111 | 56 | 6 | 39 | 0 | 9 | 421 | 7 | 389 | 0 | 397 | 11 | 402 | 0 | 0 | 0 | 412 | 0 | 438 | 0 | 275 | 100 | 1 | 0 |
| F21_18 | 1 | Kalannie | Spartacus CL | 63 | 0 | 54 | 0 | 5 | 5 | 29 | 27 | 29 | 8 | 0 | 18 | 0 | 5 | 66 | 0 | 70 | 0 | 68 | 0 | 71 | 0 | 0 | 0 | 63 | 0 | 66 | 0 | 47 | 19 | 1 | 0 |
| | 2 | | | 71 | 0 | 62 | 0 | 7 | 5 | 43 | 33 | 32 | 9 | 0 | 18 | 0 | 5 | 82 | 0 | 78 | 0 | 80 | 2 | 82 | 0 | 0 | 0 | 72 | 0 | 78 | 0 | 57 | 23 | 0 | 0 |
| F21_19 | 1 | Muresk | Spartacus CL | 537 | 0 | 541 | 0 | 16 | 0 | 487 | 54 | 73 | 36 | 10 | 6 | 21 | 7 | 557 | 0 | 568 | 0 | 587 | 0 | 557 | 0 | 0 | 0 | 535 | 0 | 578 | 0 | 222 | 326 | 10 | 0 |
| | 2 | | | 529 | 0 | 543 | 0 | 15 | 0 | 479 | 48 | 56 | 25 | 27 | 4 | 0 | 4 | 551 | 0 | 502 | 0 | 573 | 0 | 498 | 0 | 0 | 0 | 504 | 0 | 574 | 0 | 219 | 266 | 0 | 0 |
| F21_20 | 1 | Cunderdin | Spartacus CL | 250 | 12 | 267 | 10 | 16 | 0 | 269 | 24 | 24 | 6 | 0 | 17 | 0 | 17 | 251 | 22 | 279 | 0 | 188 | 78 | 280 | 2 | 0 | 0 | 282 | 0 | 280 | 0 | 125 | 163 | 3 | 11 |
| | 2 | | | 243 | 11 | 253 | 2 | 13 | 0 | 266 | 21 | 21 | 5 | 0 | 11 | 0 | 8 | 234 | 24 | 218 | 0 | 182 | 50 | 256 | 0 | 0 | 0 | 270 | 0 | 266 | 0 | 118 | 111 | 0 | 12 |

[a] The insertion position was detected as either absence or presence of the PtTi insertion with EvaGreen-based ddPCR.

[b] Probe-based assay used to specifically detect the presence of the PtTi insertion, irrespective of insertion position.

Supplementary Table S3. Fungicide groups applied in the 2021 sampling season and previous seasons as seed or in-furrow treatments and foliar applications, along with the cropping sequence from 2017 to 2021.

| Field | Location | Barley Cultivar | In-season applications[a] | | Previous Season | | Crop Sequence | | | | |
|---|---|---|---|---|---|---|---|---|---|---|---|
| | | | Seed/In-furrow | Foliar | Seed/In-furrow | Foliar | 2021 | 2020 | 2019 | 2018 | 2017 |
| F21_01 | Meckering | Spartacus CL | 4 + 11 | 3 | 7 | 3 + 11 | barley | barley | barley | wheat | wheat |
| F21_02 | Meckering | RGT Planet | 4 + 11 | Nil | 7 | 3 + 11 | barley | wheat | wheat | barley | barley |
| F21_03 | Meckering | Maximus CL | 4 + 11 | Nil | 7 | 3 + 11 | barley | wheat | wheat | lupin | barley |
| F21_04 | Doodenanning | Spartacus CL | 7 | 3 | 7 | 3 + 11 | barley | barley | barley | barley | barley |
| F21_05 | Warding East | Spartacus CL | Nil[b] | Nil | 7 | 3 | barley | barley | barley | barley | barley |
| F21_06 | Cunderdin | Spartacus CL | 7 | 3 + 11 | 7 | 3 + 11 | barley | barley | barley | barley | barley |
| F21_07 | Cunderdin | Spartacus CL | 7 | 3 + 11 | 7 | 3 + 11 | barley | barley | barley | barley | barley |
| F21_08 | Cunderdin | Spartacus CL | 3 + 4 + 7 | 3 | 7 | 3 | barley | barley | barley | barley | barley |
| F21_09 | Cunderdin | Spartacus CL | 7 | 3 | 7 | 3 | barley | barley | barley | barley | barley |
| F21_10 | Nukarni | Spartacus CL | 3 + 4 + 7 | 3 | 3 + 4 + 7 | 3 | barley | field pea | barley | barley | NA |
| F21_11 | Nukarni | Maximus CL | 3 + 4 + 7 | 3 | 3 + 4 + 7 | 3 | barley | field pea | barley | barley | NA |
| F21_12 | Nukarni | Scope CL | NA[c] | NA | NA | NA | barley | NA | NA | NA | NA |
| F21_13 | Meckering | Spartacus CL | 7 | 3 + 11 | NA | NA | barley | barley | barley | barley | barley |
| F21_14 | Quelagetting | Spartacus CL | 3 | 3 + 11 | 3 | 3 | barley | oats | barley | barley | barley |
| F21_15 | Quelagetting | Buff | 3 | 3 + 11 | 3 | 3 | barley | barley | barley | barley | barley |
| F21_16 | Watercarrin | Spartacus CL | 3 | 3 + 11 | 3 + 7 | 3 | barley | wheat | wheat | lupin | barley |
| F21_17 | Watercarrin | Spartacus CL | 3 | 3 + 11 | 3 + 7 | 3 | barley | wheat | wheat | lupin | barley |
| F21_18 | Kalannie | Spartacus CL | 3 | Nil | 3 | Nil | barley | barley | barley | barley | barley |
| F21_19 | Muresk | Spartacus CL | Nil | Nil | 3 | 3 | barley | NA | NA | NA | NA |
| F21_20 | Cunderdin | Spartacus CL | NA | NA | NA | NA | barley | NA | NA | NA | NA |

[a] FRAC Group 3 (demethylation inhibitor), Group 4 (phenylamide), Group 7 (succinate dehydrogenase inhibitor), Group 11 (quinone outside inhibitor). Numbers with '+' between indicate a mixture of fungicide groups.

[b] 'Nil' indicates no fungicide application.

[c] 'NA' indicates information was not available.

Supplementary Table S4. Mating-type copy numbers for *Pyrenophora teres* f. *teres* determined by droplet digital PCR using DNA extracted from combined net blotch lesions collected from twenty barley fields in 2021. Departure from a 1:1 ratio is indicated by the chi-square ($\chi2$) statistic.

| Field | *MAT1-1-1* | *MAT1-2-1* | $\chi^{2a}$ | *P*-value |
|---|---|---|---|---|
| F21_01 | 0 | 0 | - | - |
| F21_02 | 9 | 0 | - | - |
| F21_03 | 0 | 0 | - | - |
| F21_04 | 0 | 0 | - | - |
| F21_05 | 2 | 0 | - | - |
| F21_06 | 0 | 0 | - | - |
| F21_07 | 0 | 0 | - | - |
| F21_08 | 2 | 0 | - | - |
| F21_09 | 4 | 0 | - | - |
| F21_10 | 6 | 0 | - | - |
| F21_11 | 0 | 0 | - | - |
| F21_12 | 0 | 0 | - | - |
| F21_13 | 1 | 0 | - | - |
| F21_14 | 0 | 0 | - | - |
| F21_15 | 0 | 0 | - | - |
| F21_16 | 2 | 10 | 5.33 | 0.021 |
| F21_17 | 5 | 0 | - | - |
| F21_18 | 0 | 0 | - | - |
| F21_19 | 5 | 0 | - | - |
| F21_20 | 2 | 11 | 6.23 | 0.013 |

[a] Samples with less than 10 total mating type copy numbers did not have the chi-square ($\chi2$) statistic calculated due to a small sample size. $\chi2$ value based on one degree of freedom.

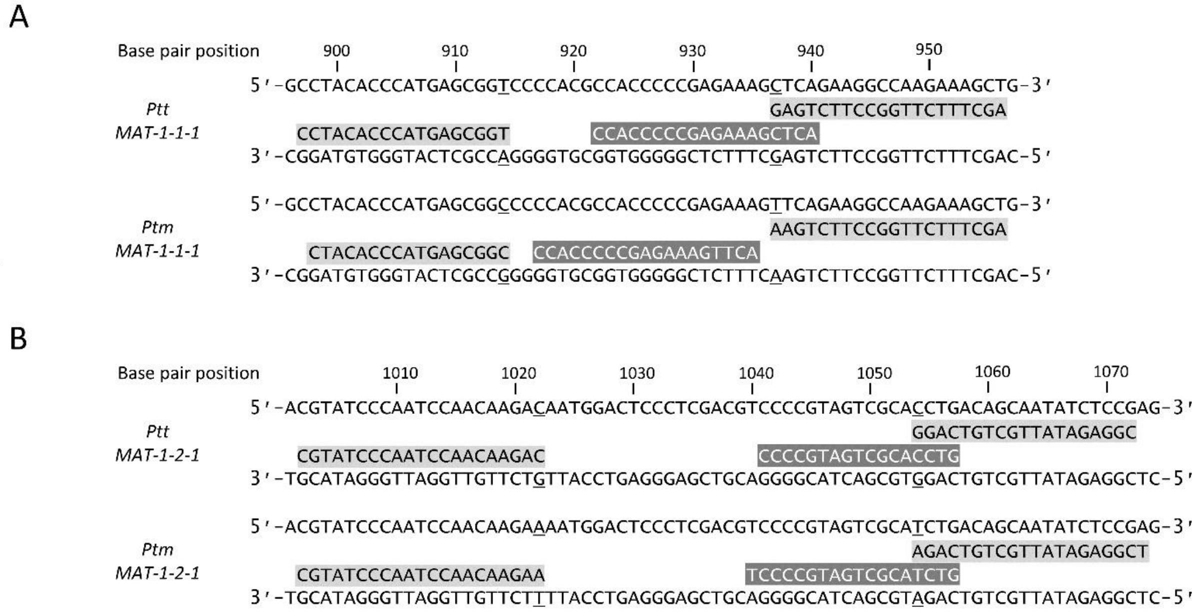

Supplementary Figure 1. Alignment of primers (black text on grey) and probes (white text on grey) against the (A) MAT-1-1-1 and (B) MAT-1-2-1 gene sequences of *Pyrenophora teres* f. *teres* (*Ptt*) *and P. teres* f. *maculata (Ptm)*. Underlined bases in the gene sequence were the single nucleotide polymorphisms targeted for specific detection. GenBank accessions for each sequence were (A) HM121991 (*Ptt*) and HM121994 (*Ptm*) and (B) HM122000 (*Ptt*) and HM122007 (*Ptm*). GenBank accession numbers (A) HM121991 and (B) HM122000 were used for base pair reference positions.
